# Treatment effects in epilepsy: a mathematical framework for understanding response over time

**DOI:** 10.1101/2024.01.22.576627

**Authors:** Gwen Harrington, Peter Kissack, John R. Terry, Wessel Woldman, Leandro Junges

## Abstract

Epilepsy is a neurological disorder characterized by recurrent seizures, affecting over 65 million people worldwide. Treatment typically commences with the use of anti-seizure medications, both mono- and poly-therapy. However more invasive therapies such as surgery, electrical stimulation and focal drug delivery may also be considered in an attempt to render the person seizure free. Although a significant portion ultimately benefit from these treatment options, treatment responses often fluctuate over time.The physiological mechanisms underlying these temporal variations are poorly understood, making prognosis one of the biggest challenges for treating epilepsy. In this work, we use a dynamic network model of seizure transition to understand how seizure propensity may vary over time as a consequence of changes in excitability. Through computer simulations, we explore the relationship between the impact of treatment on dynamic network properties and their vulnerability over time that permit a return to states of high seizure propensity. We show that, for small networks, vulnerability can be fully characterised by the size of the first transitive component (FTC). For larger networks, we find measures of network efficiency, incoherence and heterogeneity (degree variance) correlate with robustness of networks to increasing excitability. These results provide a set of potential prognostic markers for therapeutic interventions in epilepsy. Such markers could be used to support the development of personalized treatment strategies, ultimately contributing to understanding of long-term seizure freedom.

## 1 Introduction

The response to treatment in epilepsy - such as anti-seizure medication (ASM), neurostimulation, or surgery - often fluctuates over time. Most clinical studies in this context have examined this with respect to the overall long-term probability of seizure freedom for people with epilepsy (e.g. probability of no seizures ≥ 1 year) (Brodie et al. 2012; Chen et al. 2018). On shorter time-scales, perhaps the most prominent example of a transient, declining change in treatment outcomes is the so-called “honeymoon effect”. This is broadly characterized by a period of significant reduction in seizure frequency following the intervention (the “honeymoon” period), which can last from a couple of weeks to several months. By definition, the honeymoon period is followed by an eventual increase in seizure frequency, sometimes to levels at least as high as before the intervention (Boggs, Nowack, and Drinkard 2000). Understanding mechanisms that contribute to this phenomena are therefore critical for improving outcomes for people with epilepsy.

For anti-seizure medications (ASMs), the honeymoon effect (also known as acquired tolerance) is most commonly observed in benzodiazepines. As a result they are generally considered unsuitable for use as long-term treatments (Riss et al. 2008). However, people with epilepsy are known to develop tolerance to a wider range of medications (Abou-Khalil, Löscher, and Schmidt 2008; Löscher and Schmidt 2006). A study published in 2000 saw that as many of 22 out of 80 patients experienced a return to at least a baseline seizure frequency after an initial positive response to medication (Boggs, Nowack, and Drinkard 2000). Animal studies have suggested seizure type may also affect the likelihood of tolerance to certain ASMs, however little supporting evidence exists in humans (Löscher, Rundfeldt, et al. 1996; Löscher and Schmidt 2006). Further, there is the phenomenon of “cross-tolerance”. Here, tolerance to one medication may lead to tolerance to another (Abou-Khalil, Löscher, and Schmidt 2008). This observation could support the notion that tolerance emerges due to adaptation of the underlying mechanisms of seizure generation and propagation.

In the context of epilepsy surgery, patients undergoing resective surgery often experience a relapse in seizures following a successful period of remission (Petrik et al. 2021; De Tisi et al. 2011). Rates of seizure recurrence and surgical success rates are highly heterogeneous, depending both on the type of seizures experienced, the presence (or otherwise) of visible brain lesions and the nature of the surgery performed (Téllez-Zenteno, Dhar, and Wiebe 2005). Contributing factors can include incomplete resection of the epileptogenic zone, as well as the emergence of new or previously-undetected epileptogenic networks. In this latter scenario, seizures of a different nature to those experienced prior to surgery are commonplace (Petrik et al. 2021). Onset of epilepsy during the first year of life has been associated with late seizure relapse following apparently successful surgery (Petrik et al. 2021), supporting the idea that some brains may have a more ingrained tendency to ictogenicity. Other studies have also shown that seizure recurrence within 6 months of surgery is associated with people whose seizures began earlier in life (Goellner et al. 2013).

Collectively these trends in seizure propensity over extended periods of time post treatment, suggest a diversity of neurophysiological mechanisms may contribute to the return of seizures in people with epilepsy. Given the complexity of these mechanisms, as well as their diverse scales of description (e.g. spatial, temporal), a mathematical modelling approach is ideally suited to integrate these features and study how they could drive changes to seizure propensity (Goodfellow et al. 2016; Jirsa et al. 2017; Sinha et al. 2017; Junges et al. 2020). Typically these models endow regions within a network structure with a specific mechanism describing a process of interest (e.g. a transition into a seizure state). Further, the network structures are often inferred or estimated directly from imaging or neurophysiological data (Chiarion et al. 2023; Wang et al. 2014).

We have previously used dynamic network models of seizure transition to simulate and predict the effects of treatment strategies (Woldman, Cook, and Terry 2019; Junges et al. 2020). In these model frameworks, seizure propensity is fundamentally impacted by a combination of local brain excitability and network configuration. Some key aspects of dynamic brain networks (as represented in these models) are not only seizure propensity, but also robustness to change, i.e., how robust a state of low seizure propensity is to potential future changes in network reconfiguration and/or local excitability.

In this study, we systematically explore the relationship between properties of brain networks and the response of these networks to a drive towards increased seizure propensity (considered as an increase in overall excitability). These network properties range in complexity from simple measures such as the average clustering coefficient or efficiency to more complex features such as the first transitive component and trophic incoherence. They capture a variety of features of networks which may be relevant for prognosis, for example, the combined effect of network directionality and small-worldness. The drive towards increased seizure propensity represents the combined effect of the neurophysiological mechanisms that could underpin transient responses to treatment. We assess robustness to these mechanisms through quantification of the speed and extent that seizure propensity increases in these networks when a perturbation is applied, and how properties of the network impact these changes.

## 2 Materials and Methods

We consider a phenomenological model of seizure transition that permits the existence of two states, reflecting seizure-like behaviour and background activity respectively (Kalitzin, Velis, and Silva 2010; Benjamin et al. 2012). The model is a modified version of the normal form of the subcritical Hopf bifurcation, which incorporates a slowly varying time-dependent variable *λ* reflecting the level of “excitability” within a brain region. This type of model has been utilised for epilepsy in a variety of contexts (Hebbink et al. 2017; Junges et al. 2020). Within a single node, the dynamic evolution is defined by:

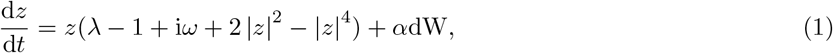

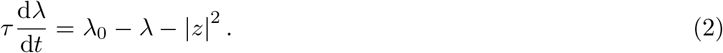

In this formulation, node activity *z* is a complex variable *x* +i*y*, such that Re(*z*) = *x* corresponds to the simulated EEG electrode activity. *ω* is the frequency of the stable limit cycle of this system, and can be tuned so that this frequency corresponds to that seen during seizure-like activity. *λ* is the time-dependent node excitability and *λ*_0_ the constant baseline level of excitability. *τ* is a real constant modulating the rate at which a node transitions from the seizure-like to the non seizure-like state. *α* reflects the impact of dynamic inputs received by regions of the brain (nodes) that are not explicitly accounted for within the model dynamics. This noise drives transitions to the limit cycle. Under an Euler-Maruyama scheme, dW draws a value from the uniform distribution bounded by 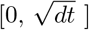 at each time-step. Typical values for these model parameters can be found within Table 1.

**Table 1:** Parameters of the modified subcritical Hopf model used in this paper (Junges et al. 2020).

| Parameter | Meaning | Value |
| --- | --- | --- |
| $N$ | Number of nodes in the network | 3, 20 |
| $\omega$ | Frequency of stable limit cycle | 20 Hz |
| $\beta$ | Diffusive coupling strength | 0-6 |
| $\gamma$ | Additive coupling strength | 0-6 |
| $\tau$ | Time-scaling factor | 5 s |
| $dt$ | Simulation time | 0.0005 s |
| $\alpha$ | Noise amplitude coefficient | 0.08 |
| $M$ | Adjacency matrix | 0, 1 |

This system is deterministic when *α* = 0. Within the physically permissible region of *λ* ∈ [0, 1] and |*z*| ≥ 0, the dynamics of *z* are characterised by three steady-state solutions. The first is a stable limit cycle at 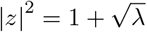. This represents seizure-like behaviour. The second is a fixed point which exists at *z* = 0 corresponding to background (inter-ictal) behaviour. Separating these two stable solutions is an unstable limit cycle at 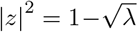. These steady state solutions are shown in Figure 1.

**Figure 1:**
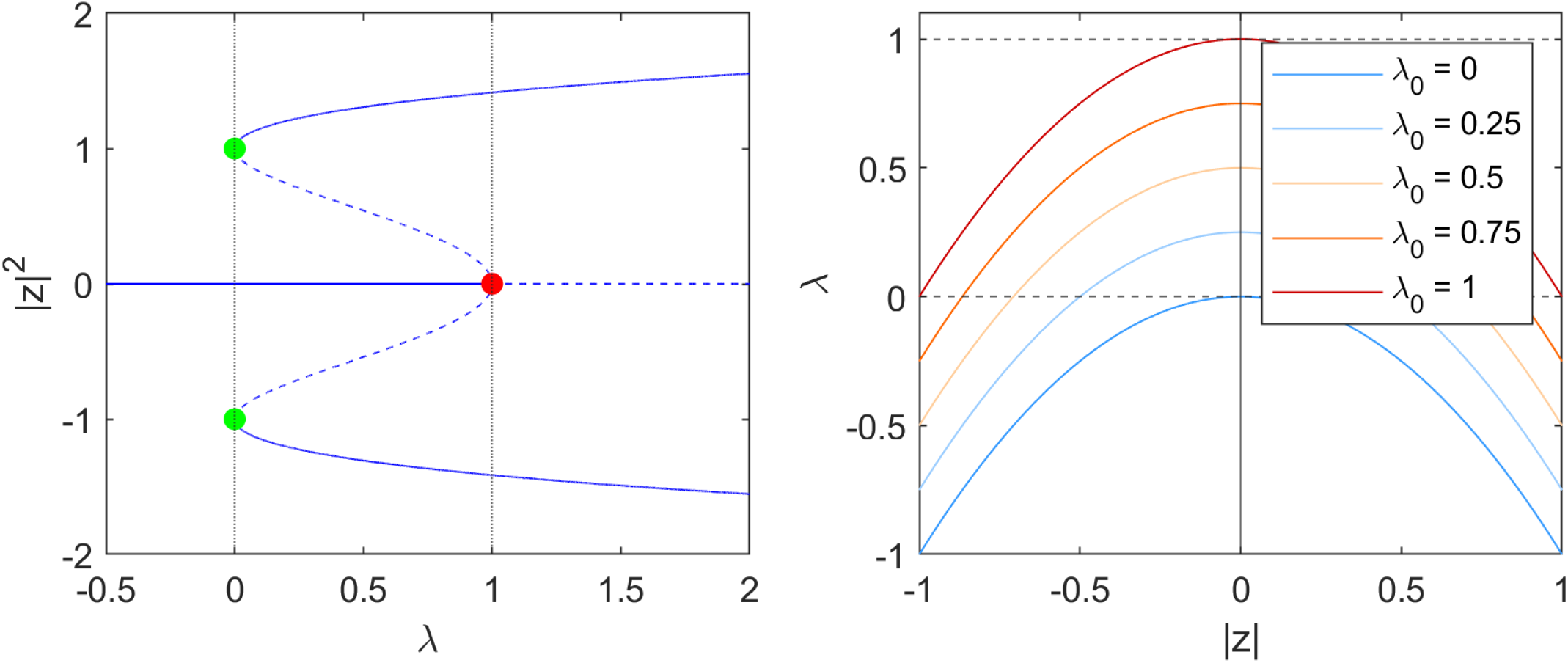
Left: bifurcation diagram of |*z*|^2^ in *λ*. Stable steady states are indicated as solid lines, and unstable steady states as dashed lines. The red point at (1, 0) represents the Hopf Bifurcation. Two limit points are marked in green at (0, 1) and (0, -1). Right: The nullclines of *λ* at different values of *λ*_0_, which is parabolic with a peak at (|*z*| = 0, *λ* = *λ*_0_). Grey dashed lines indicate the maximum and minimum values of *λ*_0_. When |*z*| = 0 in the interictal state, *λ* converges to *λ*_0_. In the ictal state, |*z*| takes on larger values and consequently *λ* converges to lower values.

Equation 2 describes the *λ* dynamics, and has a single stable steady-state solution in which *λ* = *λ*_0_ − |*z*|^2^, illustrated in Figure 1. When *z* is close to its fixed point, in the inter-ictal state, |*z*| ^2^ is very small and *λ* therefore tends to *λ*_0_; in the case where |*z*| is large, *λ* tends towards values less than *λ*_0_, decreasing the excitability and forcing a natural return of the system towards the fixed point. The time-scale of this return is determined by the slow variable *τ*. There exists a steady state in both *λ* and *z* at *z* = 0 and *λ* = *λ*_0_. As *λ*_0_ increases the size of the basin of attraction of the stable limit cycle decreases, such that the distance between the *z* = 0 and the unstable limit cycle decreases. This makes it easier for the system to undergo transition to the seizure-like state.

The dynamics in the non-deterministic system (*α >* 0) are shown for a single node in Figure 2. The system stays near the initial condition of the stable fixed point (*z* = 0, *λ* = *λ*_0_) unless the influence of the noise is strong enough that the trajectory crosses the unstable ‘separatrix’ solution causing it to approach the ictal state. Increased activity *z* results in a drop in the excitability *λ* - the key mechanism which returns the system from the seizure-like state to the inter-ictal state.

**Figure 2:**
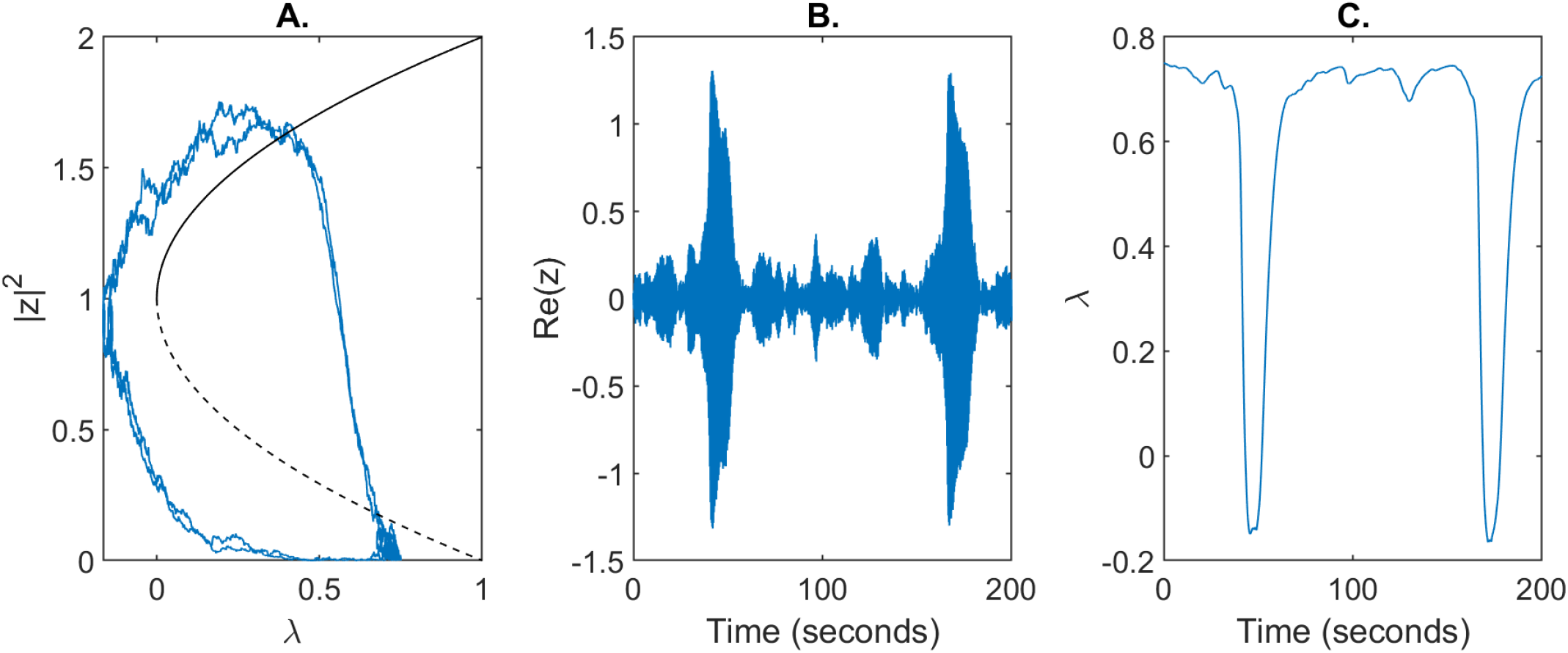
Dynamics of the modified subcritical Hopf model for a single node. A) Trajectory in phase space for a single node. The direction of flow is anti-clockwise. B) Simulated single-node EEG activity Re(z) over a timescale of T = 200s. C) Variation in ths slow excitability variable *λ*_0_.

To appropriately reflect brain network activity, we extend the model consider interacting node dynamics with a network. Here we describe network dynamics as a system of coupled time-dependent stochastic differential equations,

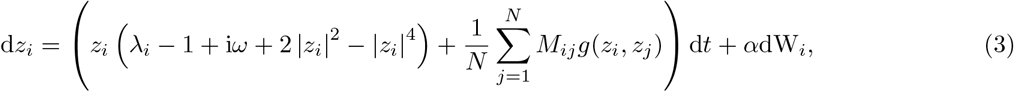

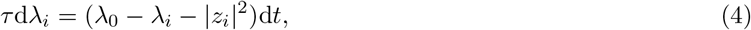

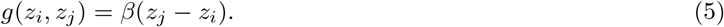

N is the total number of nodes within the network such that *i* = 1, 2, …, *N*. *M*_*ij*_ is the adjacency matrix categorising the edges within a network. We consider here directed networks without self-loops, so *M*_*ij*_ is allowed to be asymmetric with all diagonal entries set to 0. For the purposes of the present study we consider only binary networks such that entries of *M*_*ij*_ are 1 or 0, corresponding to the existence or nonexistence of an edge. *g*(*z*_*j*_, *z*_*i*_) is the diffusive coupling function where *β* is a real constant specifying the strength of the system coupling. Through this choice of coupling, interactions between nodes can additionally influence transitions from the steady-state to the stable limit cycle and vice versa.

### 2.1 Quantifying seizure propensity

In order to quantify the likelihood of the system transitioning into the seizure-like state we utilise the concept of Brain Network Ictogenicity (BNI) (Petkov et al. 2014; Lopes, Richardson, et al. 2018). Since its introduction, BNI has been used in many works as a measure of network seizure propensity. This includes as a way to quantify the change in seizure propensity after network nodes are removed or network edges are perturbed (Sinha et al. 2017; Goodfellow et al. 2016; Lopes, Richardson, et al. 2017; Lopes, Perani, et al. 2019; Lopes, Junges, et al. 2020). It evaluates the likelihood of simulated networks to transit into a seizure state, calculated as

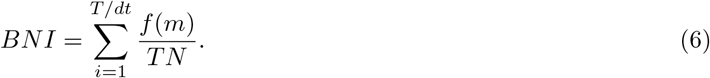

*T* is here the total simulated time-span and *dt* the time-step when system trajectories are evaluated, using a first-order Euler-Maruyama scheme. To ensure consistency between network simulations, the initial state for all nodes is the background (steady) state. *m* is the number of nodes found within the ictal state during a given time step, such that

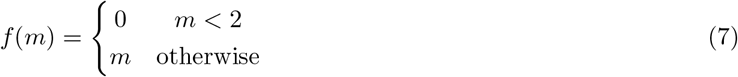

A node is considered as being in a seizure-like state if |*z*|^2^ *>* 0.5. Cases where *m <* 2 are discarded, to minimise the effect of spontaneous transition into the stable limit cycle. Consequently, if no more than 2 nodes simultaneously enter the ictal state across an entire simulation then *BNI* = 0, whereas if all network nodes remain in the high activity state throughout the simulation then *BNI* = 1. In order to exclude the possibility that any network behaviour is unique to a certain range of the coupling parameter *β*, BNI is averaged over *β* ∈ [0, 6] such that it includes behaviour from both the strong and weak coupling regimes (Junges et al. 2020). To account for the impact of stochasticity, the BNI is then re-averaged over several realisations of noise. Timescales of T = 500s or T = 2000s were chosen for each simulation with a time-step of dt = 0.0005s with 5 realisations implemented for each calculation.

In order to reproduce the increase in seizure propensity observed in the honeymoon effect, we consider a gradual increase in the baseline excitability of the network, *λ*_0_. It would be possible, in this framework, to consider higher-complexity perturbations to the system which correlate with increased seizure propensity, such as specific alterations to network topology or connection strength, however in this work we focus on excitability, the propensity for a system to attain high activity states represented by *λ*_0_, as a key feature which directly determines seizure propensity. Figure 3 shows the trajectories in BNI of two 20-node networks as a function of *λ*_0_. We consider both extremes of network connectivity - a fully-connected network, in which all possible edges are present, and an empty network, in which no edges are present. For values of *λ*_0_ *<* 0.6, BNI was uniformly 0 for both networks, such that no nodes concurrently entered the ictal state at any given point during the simulated time-span. The empty network has a much more gradual and less steep increase in BNI than the fully connected network.

**Figure 3:**
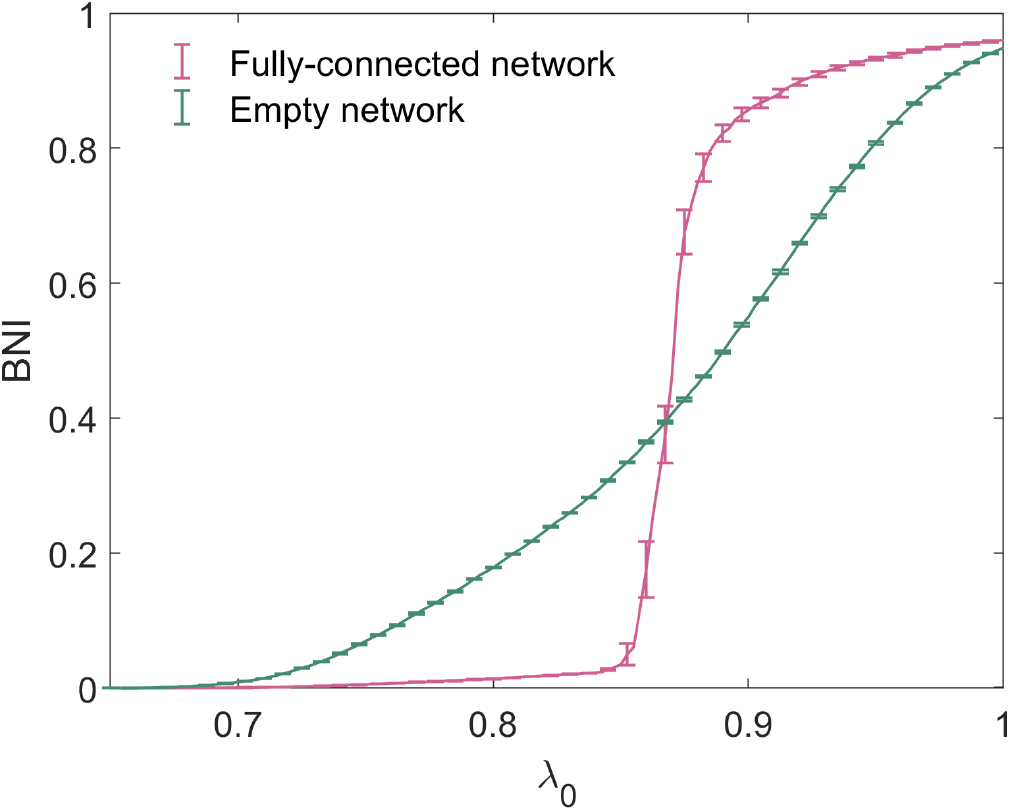
Trajectories of BNI with increasing *λ*_0_ for the fully-connected and empty 20-node networks.

We quantify the increase of network BNI under increasing *λ*_0_ by two metrics. The area under the curve (AUC) quantifies the overall increase in seizure propensity upon increase of *λ*_0_. We define the ‘quartile distance’,

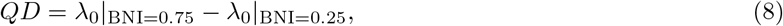

such that it is the increase in *λ*_0_ required for network BNI to increase from 0.25 to 0.75. This measure quantifies the rate of increase of network trajectories in seizure propensity as a function of *λ*_0_.

### 2.2 Network Features

In order to quantify the network characteristics associated to seizure propensity we use a number of features of binary directed networks which may have an effect on the dynamic properties of the model system. We limit our analysis to five features, though the framework proposed may be applied to any network feature that is believed to be relevant to seizure generation. We choose measures that vary in their complexity and are designed to capture a variety of network properties. We initially consider the first transitive component, a measure which identifies strongly-connected regions of the brain which may be considered “drivers” of activity, known to have a strong impact on the genesis of seizure activity (Benjamin et al. 2012). To assess the phenomenon of directionality at a more global level than FTC, we also consider the trophic incoherence, which measures the directionality of flow in the edges of a network. The presence of “driver” and “responder” regions is known to impact how different nodes and subgraphs influence each other towards similar behaviour (e.g. seizure-like or inter-ictal states) (Junges et al. 2020; Terry, Benjamin, and Richardson 2012). We further consider efficiency and mean clustering coefficient, two global measures which capture respectively global and (averaged) local characteristics of small-worldness, known to be a significant property of complex networks in nature, including of functional brain networks (Bassett and Bullmore 2006). Finally, we draw on degree variance as a measure of heterogeneity in the graph topology. Efficiency, clustering coefficient and degree variance have previously been shown to associate with epilepsy diagnosis (Faiman et al. 2021), while trophic incoherence is a relevant feature of information flow in networks (Johnson et al. 2014), making these suitable candidates for analysis of adaptations in functional brain connectivity and seizure prognosis.

#### 2.2.1 First Transitive Component

The First Transitive Component (FTC) is a measure of network connectivity introduced in the context of modelling seizures by Benjamin et al. (Benjamin et al. 2012). For a directed network of size *N* with nodes {*n*_1_, *n*_2_, …, *n*_*N*_ }, consider each distinct pair of nodes {*n*_*i*_, *n*_*j*_}. We state that *n*_*i*_ *<< n*_*j*_ if there exists a directed path from *n*_*i*_ to *n*_*j*_. Likewise, *n*_*j*_ *<< n*_*i*_ if there exists a directed path from *n*_*j*_ to *n*_*i*_. A node is a member of the FTC if every node from which it is reachable is reachable from it in return. The FTC is therefore the set of nodes *n*_*j*_ such that any *n*_*i*_ for which *n*_*i*_ *<< n*_*j*_ also satisfies *n*_*j*_ *<< n*_*i*_. It has been shown that the FTC is a predictor for the network escape times (another measure of seizure propensity which can act as a proxy for the BNI) of small networks (N − 4) (Benjamin et al. 2012). Figure 4 depicts all 13 non-isomorphic connected 3 node networks grouped by the size of FTC, which we denote as *n*.

**Figure 4:**
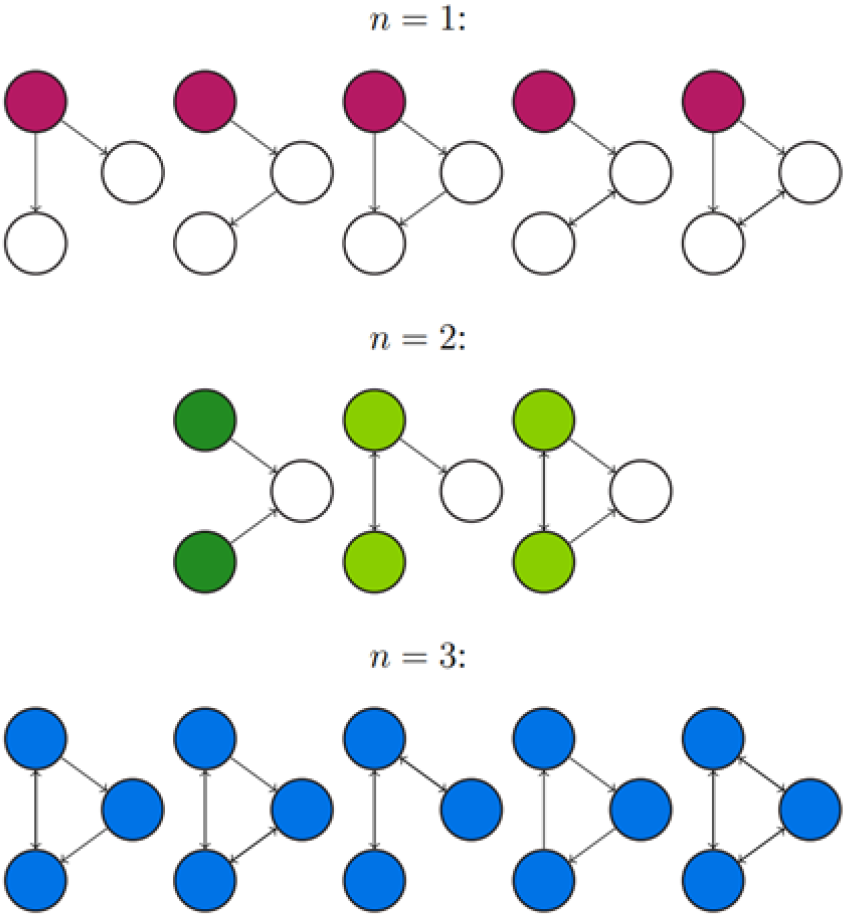
All connected directed networks of size 3 and their first transitive components. FTCs of size 1, 2, and 3 are highlighted in purple, green and blue respectively. The first network for *n* = 2 is coloured a different green to highlight that the FTC comprises two separate components of the graph, which are not connected by an edge. This leads to distinct behaviour from other networks with FTC of size 2.

#### 2.2.2 Trophic Incoherence

Trophic incoherence is a quantification of the extent to which the flow of information in a directed network is non-hierarchical (Johnson et al. 2014). The revised notion of trophic incoherence used here was introduced by MacKay et al (MacKay, Johnson, and Sansom 2020). Since the networks under consideration are unweighted, we will consider the definition in the case where all edges are assigned weight 1.

In the uniformly-weighted case, the imbalance of a node is

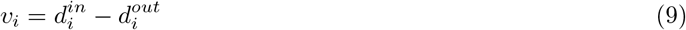

and the total weight of the node is

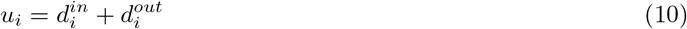

The trophic level is then defined as the vector solution *h* to the system of equations

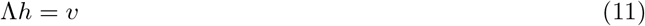

where Λ is the graph Laplacian operator

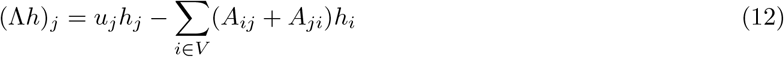

Finally, then, the trophic incoherence for a uniformly-weighted graph is

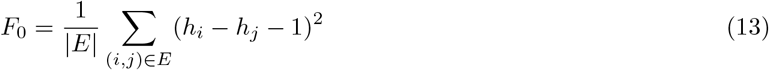

#### 2.2.3 Efficiency

The (global) efficiency (Latora and Marchiori 2001) of a directed network is defined as

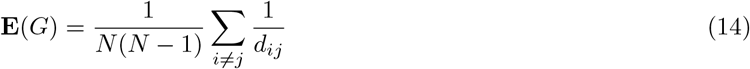

where *d*_*ij*_ is the shortest path length from node *i* to node *j*, and its inverse thus describes the efficiency of communication between the two nodes. The global efficiency is therefore an average of efficiencies between all (ordered) pairs of nodes in the directed network. Efficiency is an important marker in functional brain connectivity networks, as it represents the ability of regions of the brain to share information, and describes the global behaviour of small-world networks (Rubinov and Sporns 2010; Latora and Marchiori 2001).

#### 2.2.4 Mean Local Clustering Coefficient

For a binary directed network, the notion of mean clustering coefficient is defined as follows (Fagiolo 2007):

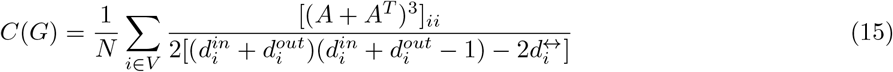

Here, 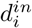 and 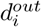 refer to the in-degree and out-degree of node *i*, and 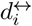 refers to the number of nodes to which *i* is connected by an edge in both directions. *A* is the adjacency matrix of the network *G*, such that [(*A* + *A*^*T*^)^3^]_*ii*_ is the number of triangles that node *i* is included in, irrespective of the directions of the edges.

Clustering measures the prevalence of small, well-connected cliques in the graph, and is a measure of the local behaviour of small-world networks (Watts and Strogatz 1998).

#### 2.2.5 Degree Variance

The variance of the node degrees quantifies the heterogeneity of a graph (Snijders 1981). For directed networks, the degree in question is taken to be the out-degree, such that the degree variance is given as

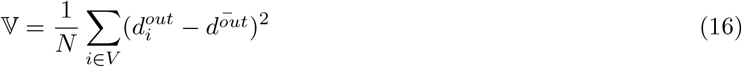

where 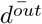 is the average out-degree of all nodes, equalling 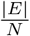, where *E* is the set of all directed edges in the network.

### 2.3 Selection of Networks

A set of 10,000 20-node random binary directed networks with a mean degree of 2.5 were generated for analysis within this paper. This network size and mean degree is in line with typical functional connectivity networks obtained from scalp or intracranial EEG recordings (Aminoff 2012; Sargolzaei et al. 2015; Lopes, Richardson, et al. 2017). Networks were generated using the NetworkX package in Python (Hagberg, Swart, and S Chult 2008), which utilises the Erdös-Rényi algorithm (Erdös and Rényi 1959; Gilbert 1959) and specified as at least weakly connected. Figure 5 shows the distribution of the number of nodes contained in the FTC, *n*, for these networks. We observe a bi-modal distribution, where the majority of networks lie in the range of *n <* 5 or *n >* 15. An equivalent distribution for 1000 networks generated with mean degrees of 1.5 and 6 is shown for comparison. As the mean degree of generated networks increases, so does the likelihood of the FTC spanning the entire network. Equivalently as networks become more sparse, the likelihood that a large proportion of the network is contained within the FTC decreases. Our choice of a mean degree of 2.5 is a pragmatic choice, such that it contains a large sample of networks with both high and low *n*. BNI has been shown to depend on the network mean degree (Petkov et al. 2014). In order to avoid such influence, networks were generated with both equal mean degree and number of nodes N.

**Figure 5:**
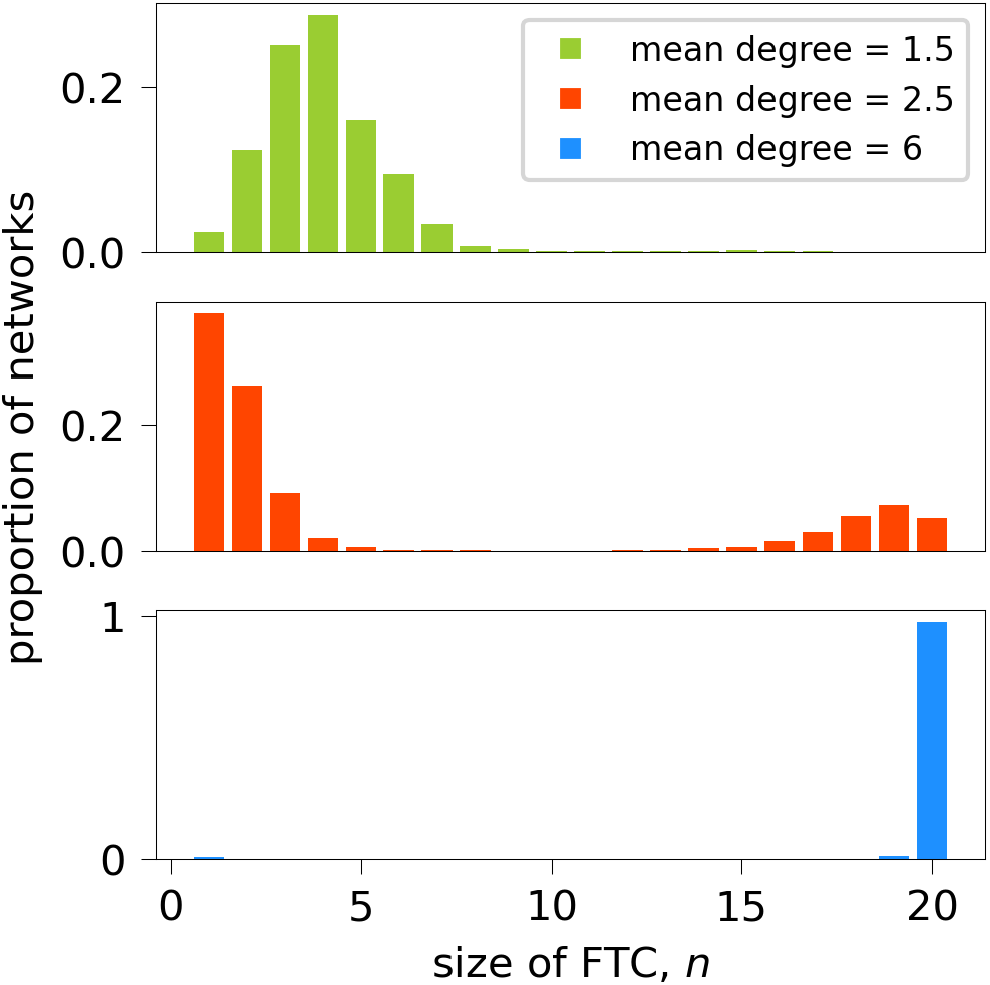
Sizes of FTCs of 10,000 randomly-generated 20-node networks of mean degree 2.5, and 1000 networks of mean degree 1.5 and 6. In the case of mean degree 2.5, for each *n* between 6 and 14, there were fewer than 50 networks (*<* 0.5%) in the sample. The least-represented size of FTC was *n* = 9, which appeared twice in the sample of 10,000. For the sample of 1000 networks with mean degree 1.5, the number of networks for each *n* ≥ 9 was less than 5, with no networks having *n* ≥ 18. For the sample of 1000 networks with mean degree 6, there were no networks with 2 ≤ *n* ≤ 18, and 97.5% of the networks had *n* = 20.

### 2.4 Correlations

Nonlinear correlations are used to measure the relationships between network measures and measures of the increase in BNI with increasing *λ*_0_. We calculate Kendall correlation coefficients between AUC/QD and efficiency, mean local clustering coefficient, trophic incoherence and degree variance; both across all sizes of FTC and for individual sizes of FTC. Significance is evaluated at *α* = 0.0001, with a Bonferroni correction for multiple comparisons.

## 3 Results

### 3.1 3 node networks

In Figure 6 we present the BNI for all 13 non-isomorphic 3 node networks (Benjamin et al. 2012) as a function of the baseline excitability *λ*_0_. We observe that network trajectories in *λ*_0_ show a tendency to group together according to the size of their FTC, *n*. This is consistent with the work of Benjamin et al. (Benjamin et al. 2012) who have shown that network escape times for 3 node networks group according to *n*. This behaviour cannot be trivially approximated by counting the number of edges within each network, e.g. 3-edge networks with *n* = 1, 2, or 3 exhibit distinct behaviour. The set of networks with *n* = 1 exhibit the most gradual increase in BNI as *λ*_0_ increases. The initial incline of networks with *n* = 2 is intermediate (between *n* = 1 and *n* = 3), with a sharp increase in incline occurring in the region of *λ*_0_ = 0.85. Networks with *n* = 3 exhibit similar behaviour, albeit with a steeper increase in BNI. The exception to this pattern is the 3-node network consisting of a single sink driven by two otherwise isolated nodes (*n* = 2), indicated in Figures 4 and 6.

**Figure 6:**
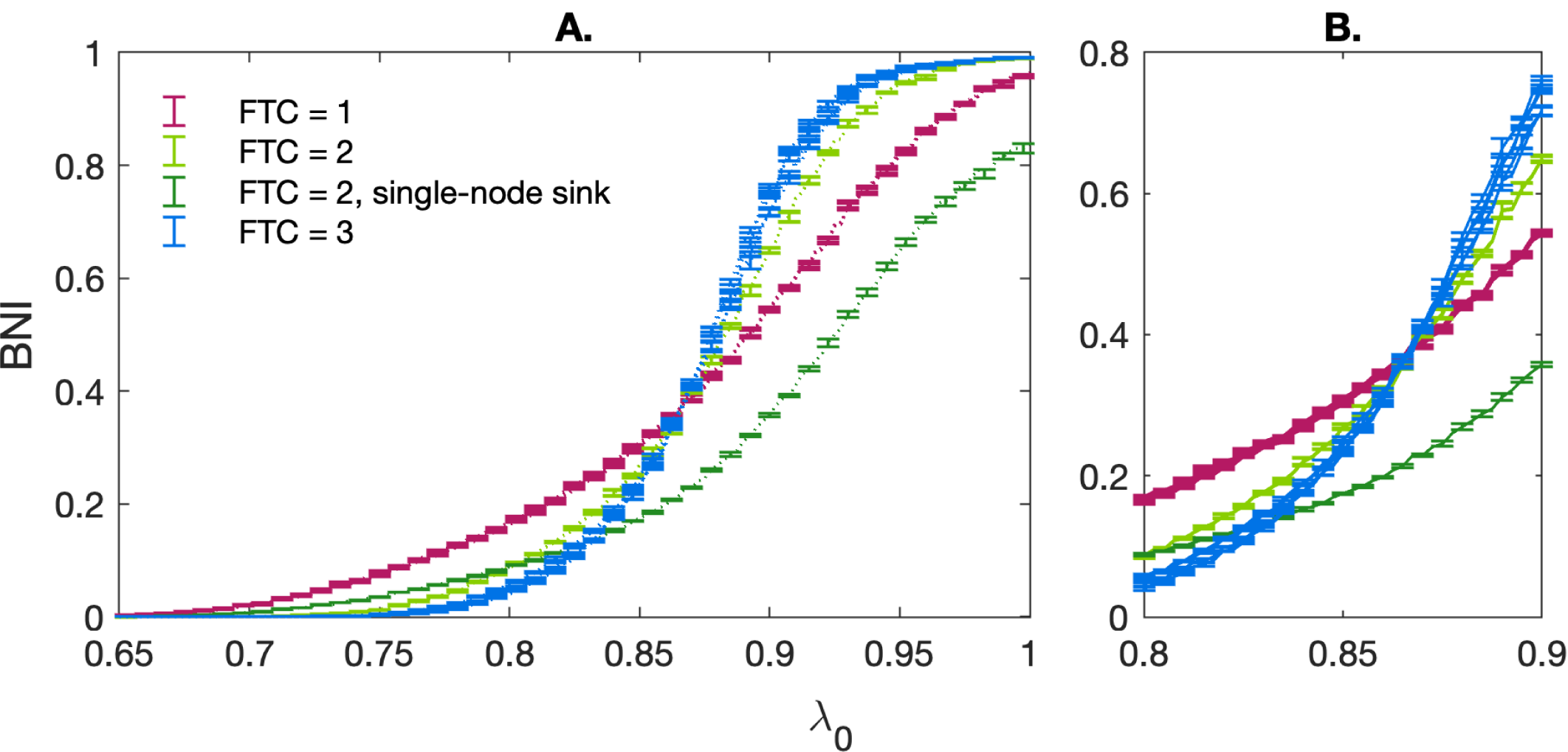
A) BNI as a function of increasing baseline excitability *λ*_0_ for all non-isomorphic three-node networks. All parameters follow the standard values stated in Table 1. Networks with an FTC containing 1 node are shown in purple, those with an FTC containing 2 nodes shown in green and those with an FTC containing the whole networks are shown in blue. The trajectory for a single-node sink (ref Fig 4) driven by two otherwise isolated nodes (*n* = 2) is shown in dark green. Errors are calculated as the standard deviation over 5 realisations of the noise co-efficient *α*. B) Focus on the region of 0.8 *< λ*_0_ *<* 0.9 where trajectories intersect.

The behaviour of these network trajectories was quantified using two measures - the area under the curve (AUC) and the quartile distance (QD). The distribution of these measures for *n* = 1, 2, 3 is shown in Figure 7. When the single-node sink is excluded, it is clear that the distribution of AUC/QD is dissimilar for different values of *n*.

**Figure 7:**
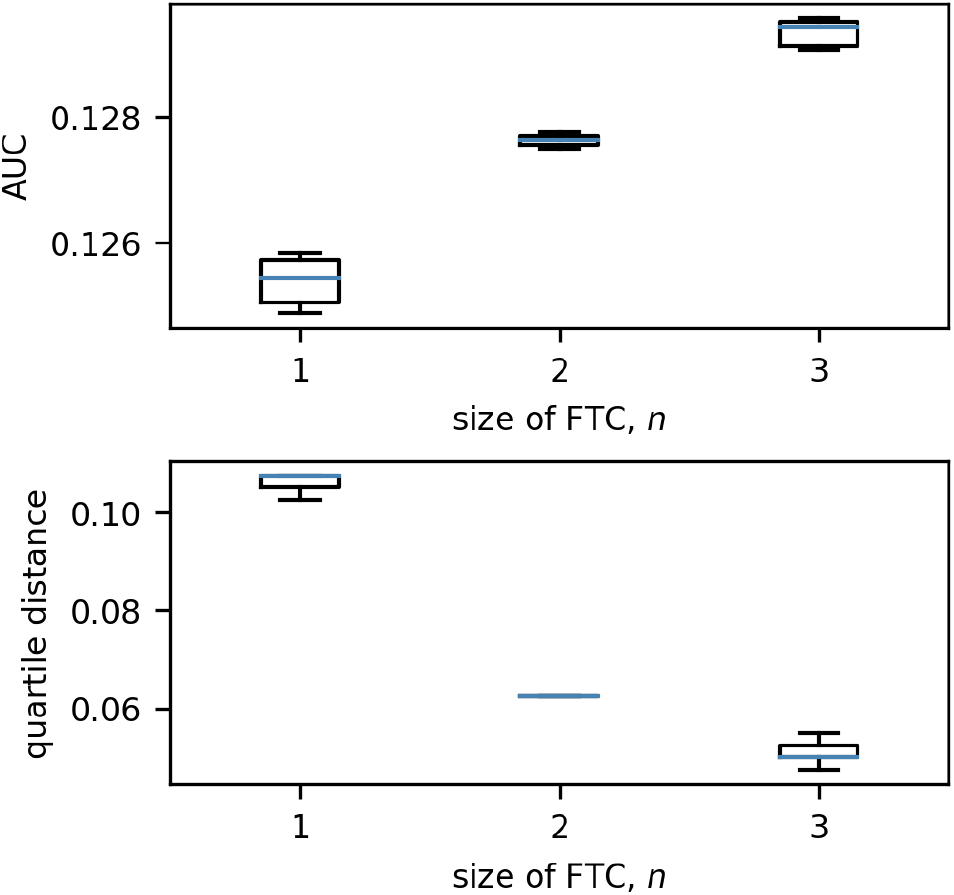
Boxplots of the distributions of quartile distance and AUC for each size of FTC in 3-node networks. We exclude the anomalous case for *n* = 2 in which the two nodes in the FTC are not connected by an edge (see Figures 4 and 6; this network has an AUC of 0.0904 and QD of 0.0950). We see that there are clear trends exhibited in the remaining: as *n* increases, AUC increases and QD decreases. Boxes are drawn from the first to third quartiles with the median value marked in blue. Whiskers are drawn at the maximum and minimum data points of the set, excluding any outliers. Outliers are marked by asterisks (*).

### 3.2 20 node networks

Having characterised the dynamics in small networks (*N* = 3), we now consider networks of size *N* = 20. Figure 8 presents the distribution of QD/AUC measures for increasing values of *n*. For both measures, values tend to remain similar for networks with FTC sizes beyond *n* = 6.

**Figure 8:**
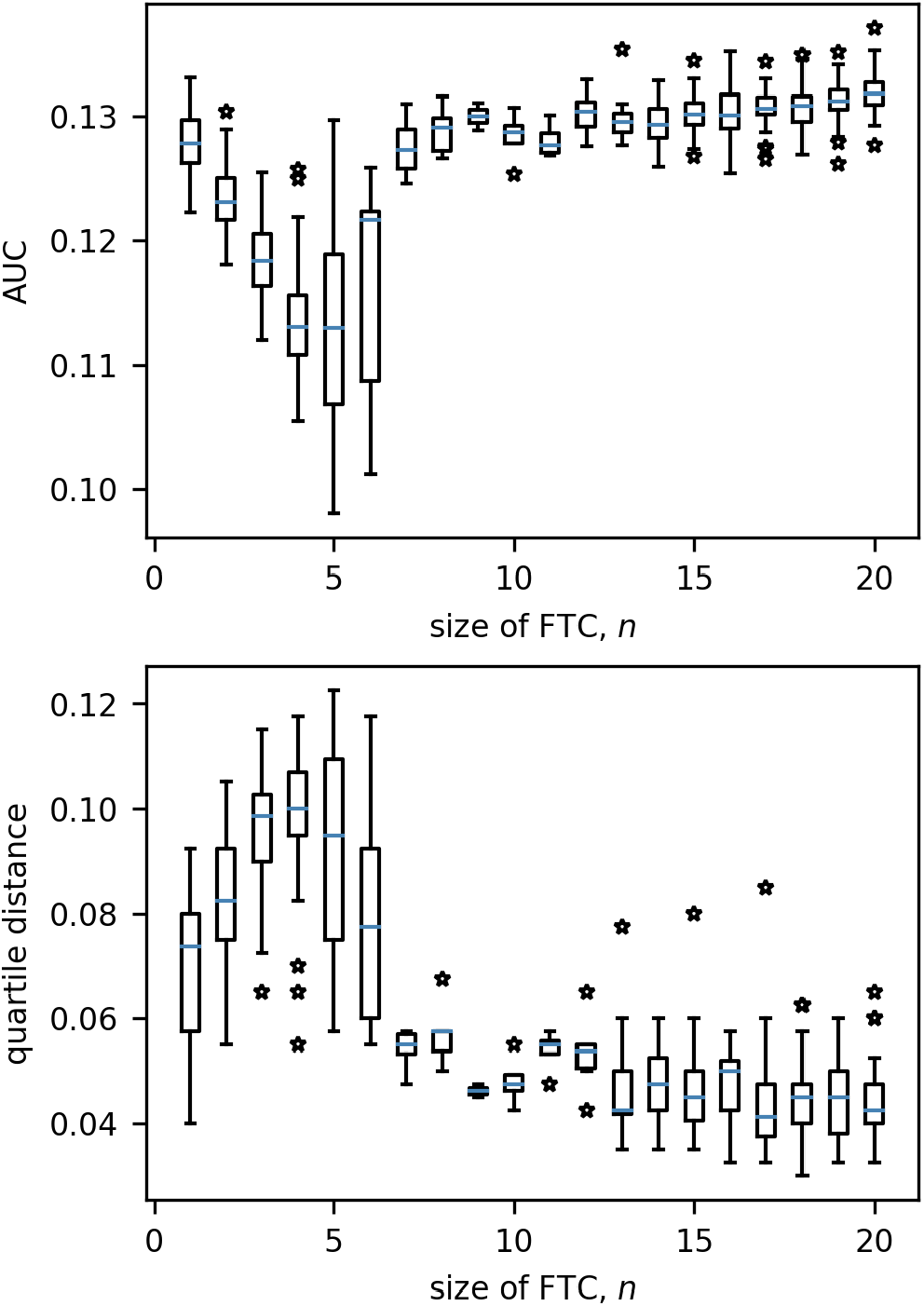
Boxplots of the distributions of quartile distance and AUC for each size of FTC in 20-node networks. Beyond *n* = 6 we see very little variation in either measure which can be associated to *n*. Boxes are drawn from the first to third quartiles with the median value marked in blue. Whiskers are drawn at the maximum and minimum data points of the set, excluding any outliers. Outliers are marked by stars.

As *n* alone is no longer sufficient to predict how network BNI will evolve as *λ*_0_ increases, we quantified the relationship between additional network measures and AUC/QD. It should be noted that the number of networks in the range 6 ≤ *n* ≤ 14 was very low (*<* 50 for each *n* out of a total 10,000, while *>* 2000 are present for *n* = 1, 2). The networks within this range were combined with a sample of 50 networks for each remaining value of *n*, and evaluated by efficiency, mean clustering coefficient, trophic incoherence and degree variance.

Figure 9 displays the relationships of network measures with AUC and QD across all sizes of FTC. Significant positive correlations were observed for AUC with network efficiency (Kendall tau = 0.393, *p <* 0.0001) and trophic incoherence (Kendall tau = 0.357, *p <* 0.0001). A significant negative correlation was observed for AUC with degree variance, (Kendall tau = -0.472, *p <* 0.0001). No strong correlation was found between AUC and mean clustering coefficient (Kendall tau = -0.0827, *p* = 1.670 *×* 10^−3^). Significant negative correlations were observed for QD with network efficiency (Kendall tau = -0.347, *p <* 0.0001) and trophic incoherence (Kendall tau = -0.398, *p <* 0.0001). A significant positive correlation was observed for QD with degree variance, (Kendall tau = 0.567, *p <* 0.0001). No significant correlation was found between QD and mean clustering coefficient (Kendall tau = 0.0575, *p* = 0.03153).

**Figure 9:**
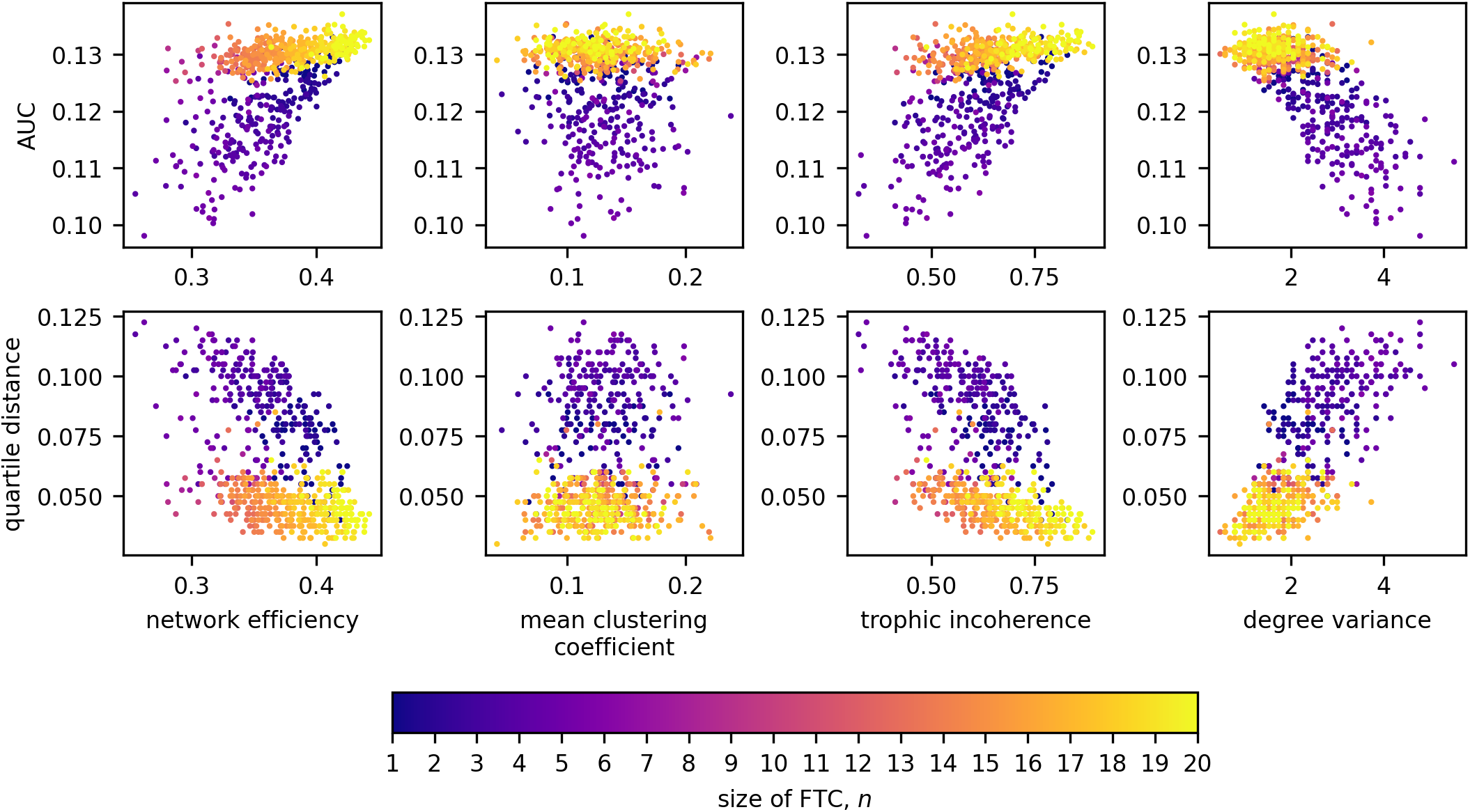
Scatter plots detailing the relationships between AUC and network measures, coloured by the size of FTC.

We will now observe in closer detail these relationships for each size of FTC. Due to the low numbers of networks generated for the intermediate values of *n*, further analysis is conducted only on high and low sizes of FTC: 1 ≤ *n* ≤ 5 and 16 ≤ *n* ≤ 20. For these, Kendall correlation coefficients between our network metrics, and network AUC/QD are shown in Figures 10.

**Figure 10:**
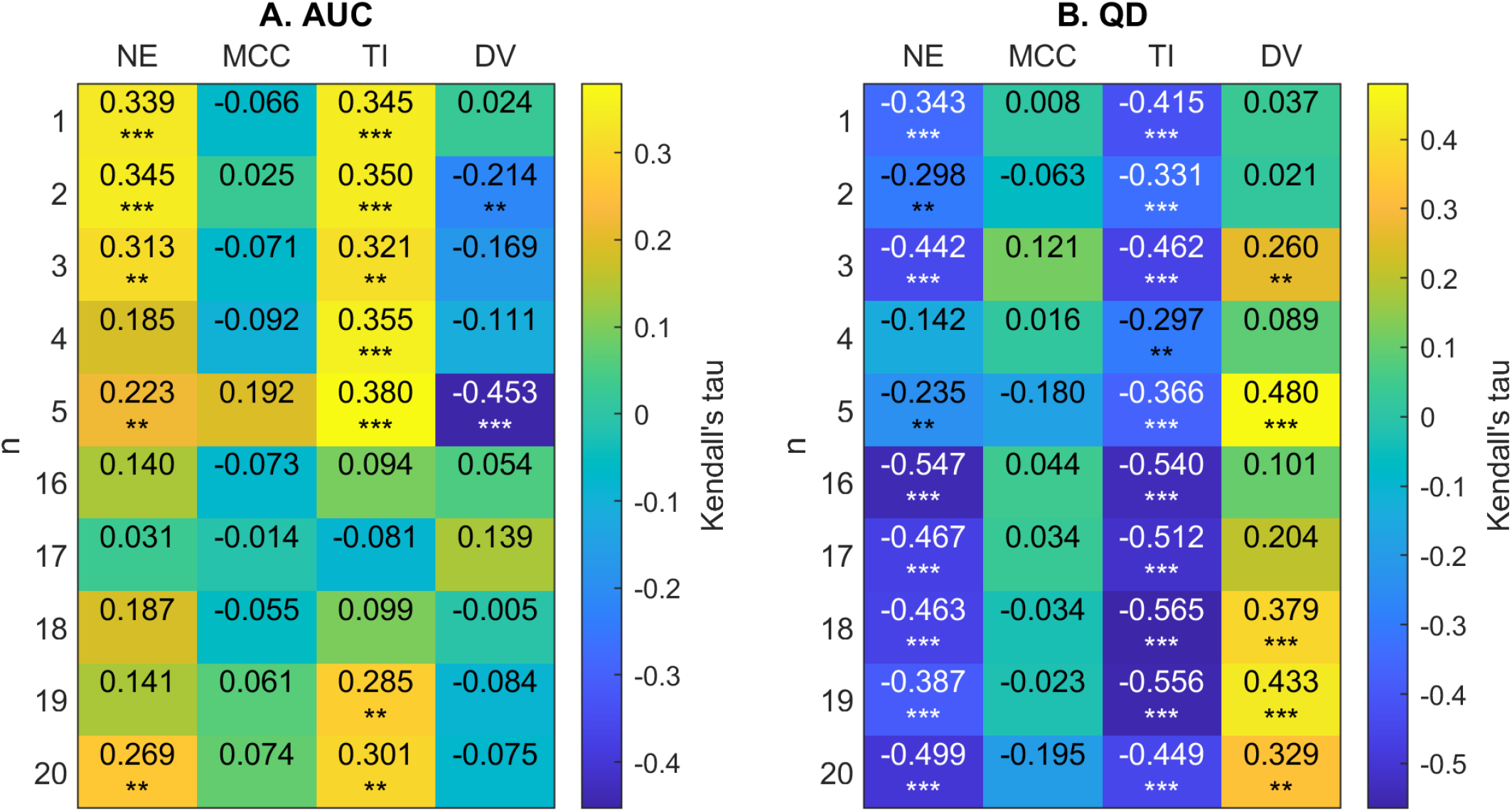
A) 2D correlation plot between AUC and network metrics (efficiency (NE), mean clustering co-efficient (MCC), trophic incoherence (TI) and degree variance (DV)) for *n* = 1 − 5, 16 − 20. Kendall correlation coefficients are colour-coded under the right axis colorbar with exact values shown in each cell. Significance values are labelled as * where *p <* 0.05, ** where *p <* 0.01 and *** where *p <* 0.0001. B) Correlation plot between QD and network metrics.

Figure 10 shows that there is a clear trend for a negative correlation between the network efficiency and QD, and between trophic incoherence and QD. For high values of *n*, there is a weak positive correlation between degree variance and QD. No consistent trend in correlation is shown between QD or AUC and mean clustering co-efficient for high or low *n*. For networks with *n* ≤ 5 there is a trend of moderate positive correlation of AUC with trophic incoherence and network efficiency, respectively.

To illustrate the results shown above, the trajectories of highest and lowest efficiency and trophic incoherence for *n* = 3, 18 are shown in Figure 11. For both *n* = 3 and *n* = 18, the initial incline of the high-efficiency network is lower than that of the low-efficiency network. The maximum slope of the high-efficiency networks is greater, with trajectories intersecting at approximately *λ* = 0.85. Similar behaviour is shown for the networks with highest and lowest trophic incoherence. Equivalent plots for *n* = 1, 5, 16, 20 can be found in Figs. S1-S4 in the Supplemental Data.

**Figure 11:**
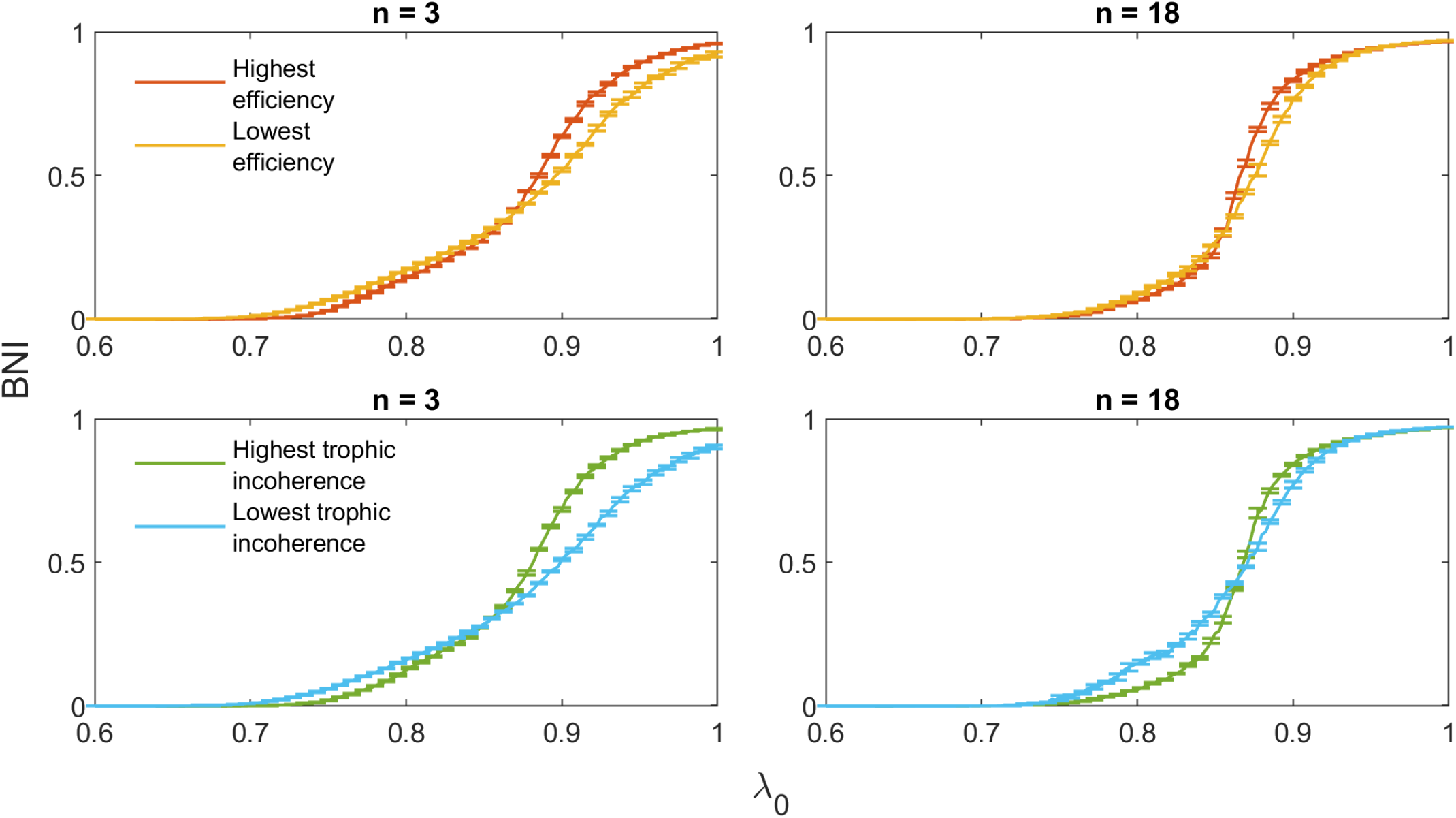
Comparison of trajectories with minimal and maximal values of efficiency and trophic incoherence for *n* = 3, 18.

It is important to point out that the relationship between network measures and AUC/QD can be influenced by the choice of network mean-degree (see Fig. S5 in the Supplemental Data). However, regarding practical applications, this is not a limiting factor of this framework as connectivity networks obtained from brain imaging can be thresholded to match the values presented in this work.

## 4 Discussion

In this study, we propose a model framework that can be used to understand changes in treatment response over time. This framework considers the interaction of a set of network features: size of the first transitive component (FTC), efficiency, incoherence, and heterogeneity (degree variance).

Our results show that, for small networks (3 nodes), distinct patterns of BNI increase as a function of the baseline excitability (*λ*_0_) can be observed. These patterns were shown to be well characterised by the number of nodes in the FTC. Networks with FTC = 1 have the most gradual (less steep) increase in BNI. From the nature of these networks (seen on the first row in Fig. 4), it is observed that the dynamics are mainly influenced by a source node, shown in purple in all networks. This node is not being “controlled” (i.e., forced to keep in the fixed point via diffusive coupling) by any other node, therefore the transition into seizure states is mainly dictated by the effect of the smooth increase in *λ*_0_ on the driving node.

For networks with FTC = 3, the graph is strongly connected, and all nodes are controlling each other to some extent. This explains why these networks remain with low values of BNI when 0 *< λ*_0_ ⪅ 0.85. However, as *λ*_0_ increases, once nodes start to transit into the limit-cycle they encourage other nodes to do the same, and the high levels of synchronization in these networks lead to a steeper increase in BNI.

For networks with FTC = 2, levels of BNI increase are intermediary when two interconnected nodes are influencing a third node (light green in Fig. 4). In this scenario, we observe a similar but less strong effect than what is seen for networks with FTC = 3 (fully connected). However, when two sources are independently influencing a third node (dark green in Fig. 4), the profile of BNI increase is much slower than for any other networks. In this case, the competing sources struggle to arrest the third node at any given time, and the values of BNI are comparatively small even for large values of *λ*_0_ (see Fig. 6). The results observed here are in line with the analysis of 3- and 4-nodes networks presented in (Junges et al. 2020). The different behaviour observed when competing effects are present in the network suggests a more complex relationship between BNI and *λ*_0_ for larger networks. Effectively this occurs, due to sub-networks that interact in intricate ways.

We see this phenomena clearly for larger networks (20 nodes), where the size of the FTC is not enough to fully characterize the relationship between BNI and *λ*_0_. Figure 8 shows that for small FTC sizes (*n* ⪅ 5), a negative (positive) correlation is observed between *n* and AUC (QD), with the lowest values of AUC (highest QD) being observed for FTC sizes of 4-5. Networks with larger FTC sizes (*n >* 6) tend to present increased AUC and decreased QD. A more detailed investigation shows that AUC tends to be higher and QD tends to be lower in more efficient, incoherent and heterogeneous networks (Fig. 10). This relationship with AUC is clearer in networks with small FTC. Interestingly, the clustering coefficient does not seem to influence a network’s response to an increase in *λ*_0_.

The results described above suggest that optimal robustness to an increase in baseline excitability are observed for networks with FTC sizes of 4-5. Minimal AUC and maximal QD suggest that increases in the baseline excitability have a smaller and more gradual effect in seizure propensity in these networks. At the same time, a similar robustness effect is seen for networks with lower efficiency, incoherence and higher degree variance.

Previous works compared functional networks obtained using electroencephalography (EEG) recordings from people with epilepsy and healthy controls. A recent literature review of biomarkers of idiopathic epilepsy from resting-state EEG explored graph-based markers and showed evidence that, in specific frequency bands, networks from people with epilepsy tend to have elevated mean degree, degree variance, and path length (Faiman et al. 2021). The observed effect of epilepsy in these measures are in line with the results presented in this work. However it is important to notice that this study explores the relationship between the network markers and the honeymoon effect, and not with seizure propensity itself. The relationship between trophic incoherence and epilepsy, to the best of our knowledge, has not been explored. Nevertheless, studies have shown that spreading processes in networks (such as seizures) can be strongly affected by trophic organisation (Klaise and Johnson 2016).

Once validated using networks obtained from brain imaging modalities (e.g., electroencephalography) and long term post-intervention seizure monitoring, the framework presented in this work can provide prognostic markers of seizure propensity progression. This will support the development of personalized intervention strategies, aiming to achieve long term seizure freedom. In the context of surgical intervention, network-based models of seizure transition have been extensively used to predict optimal surgical strategies via the estimation of seizure propensity after intervention (Goodfellow et al. 2016; Lopes, Richardson, et al. 2017; Sinha et al. 2017; Lopes, Junges, et al. 2020). However, these methods tend to ignore potential long-term changes in the system, which could lead to seizure recurrence. The framework presented in this study can significantly extend the current potential of these models by quantifying robustness to mechanisms associated to the honeymoon effect. This would ultimately contribute to support pre-surgical planning via the identification of strategies leading to more robust seizure control.

A limitation of this work is the relatively simple description of the mechanisms leading to the increase in seizure propensity after therapeutic interventions (honeymoon effect). Here, we are representing such effect via a linear and spatially homogeneous increase in baseline excitability. In reality, such effect might be best represented by complex combinations of local and more intricate changes in excitability, as well as in the network structure itself. Additionally, the specific nature of these mechanisms can depend on several factors, like epilepsy type, age at intervention (neurodevelopmental stage), and/or intervention type. However, this work does not aim to propose a tool to comprehensively estimate the effects of increased seizure propensity in every circumstance. Instead, we propose a general framework to represent the honeymoon effect, which can be modified and validated to describe specific scenarios and provide valuable prognostic markers.

## Author Contributions

GH and PK: Methodology. All authors: conception and design, analysis and interpretation, drafting and revising the manuscript.

