## Supplemental Figures S1-S5 for "Treatment effects in epilepsy: a mathematical framework for understanding response over time"

Supplementary Material

### 1 Supplementary Tables and Figures

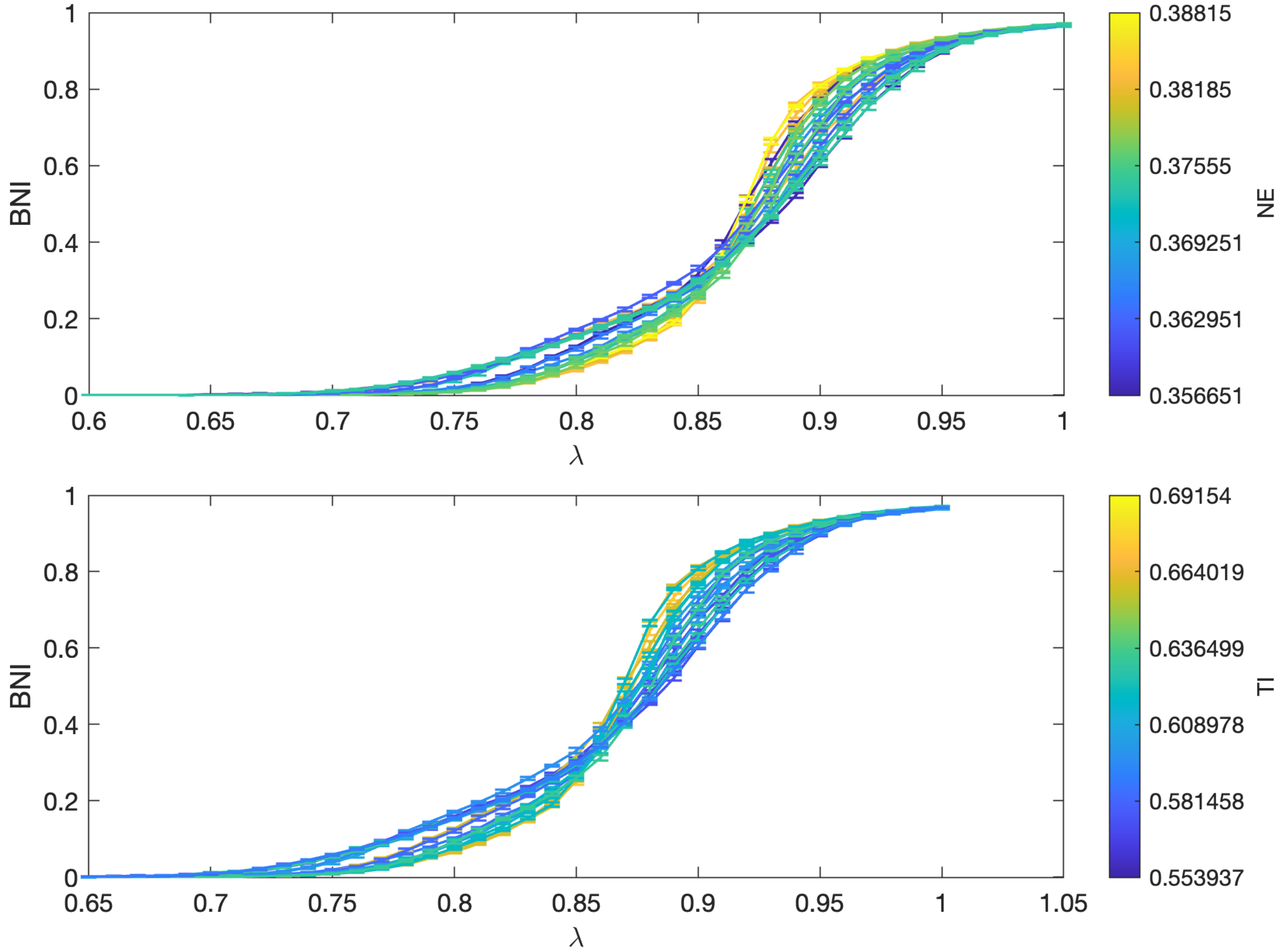


Figure S1: Increase of BNI as a function of $\lambda_0$ for 20 node networks with size of FTC, $n = 1$ and a mean degree of 2.5. Network trajectories are coloured according to their efficiency (top) and trophic incoherence (bottom).

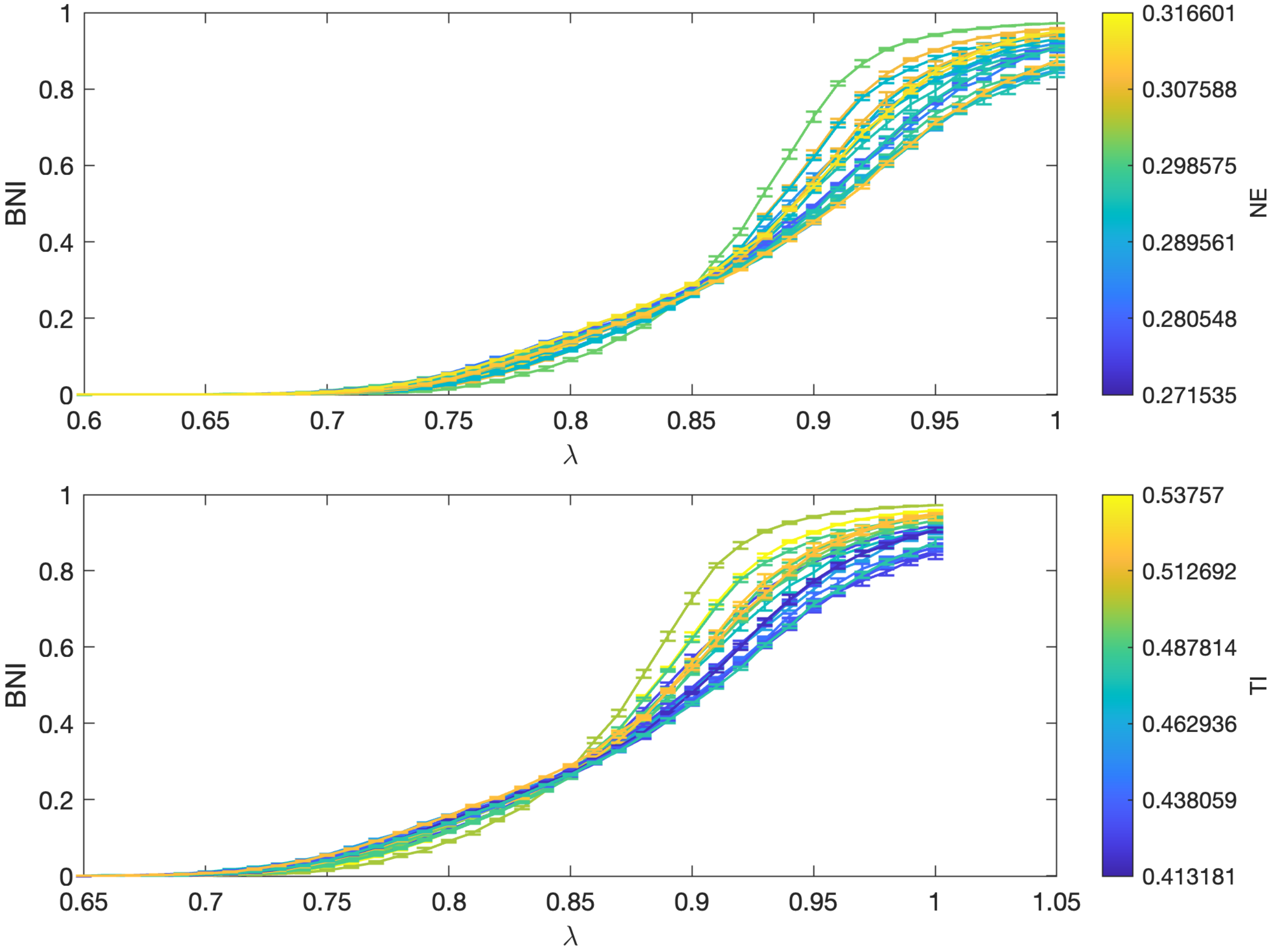


Figure S2: Increase of BNI as a function of $\lambda_0$ for 20 node networks with size of FTC, $n = 5$ and a mean degree of 2.5. Network trajectories are coloured according to their efficiency (top) and trophic incoherence (bottom).

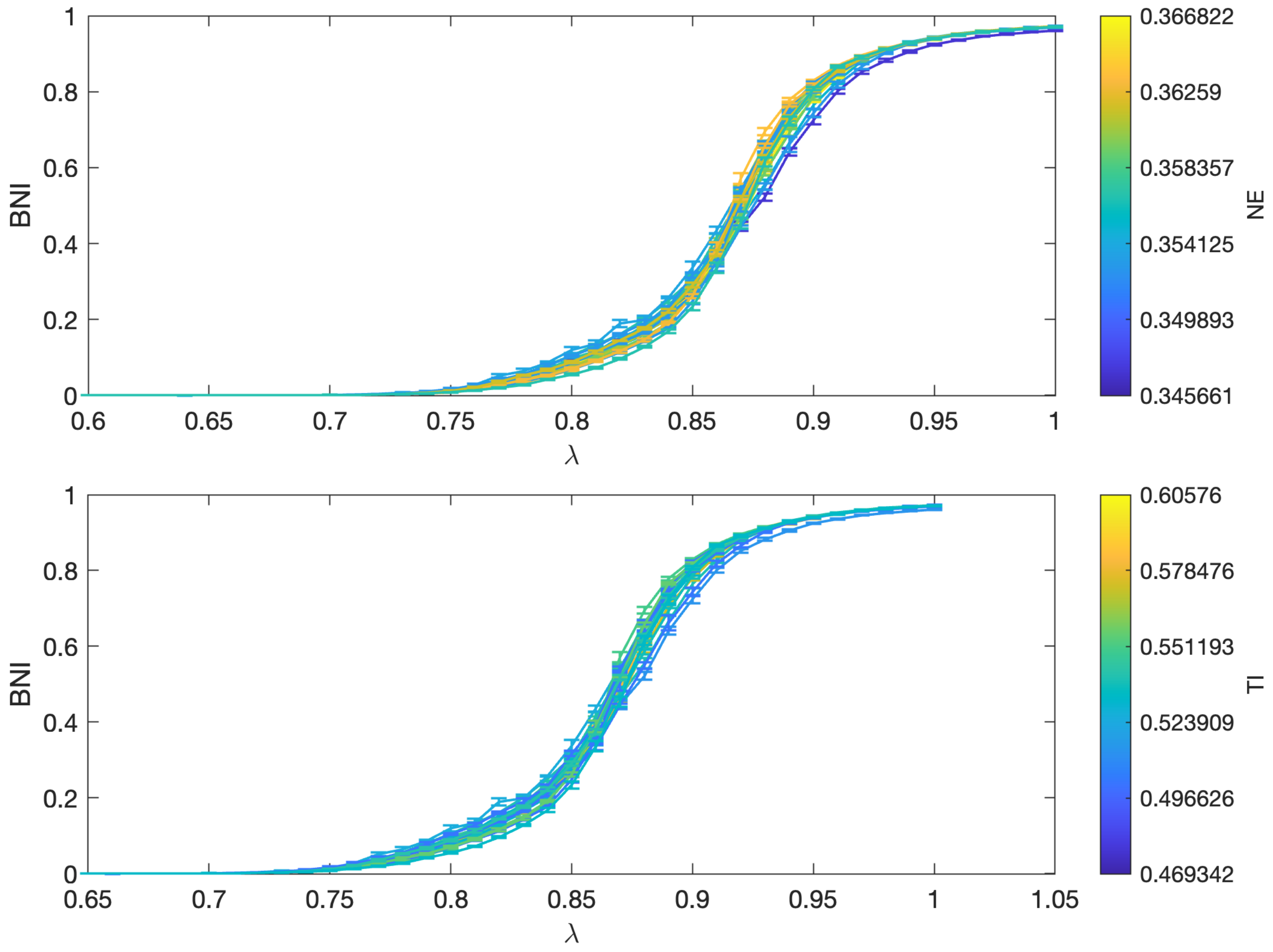


Figure S3: Increase of BNI as a function of $\lambda_0$ for 20 node networks with size of FTC, $n = 16$ and a mean degree of 2.5. Network trajectories are coloured according to their efficiency (top) and trophic incoherence (bottom).

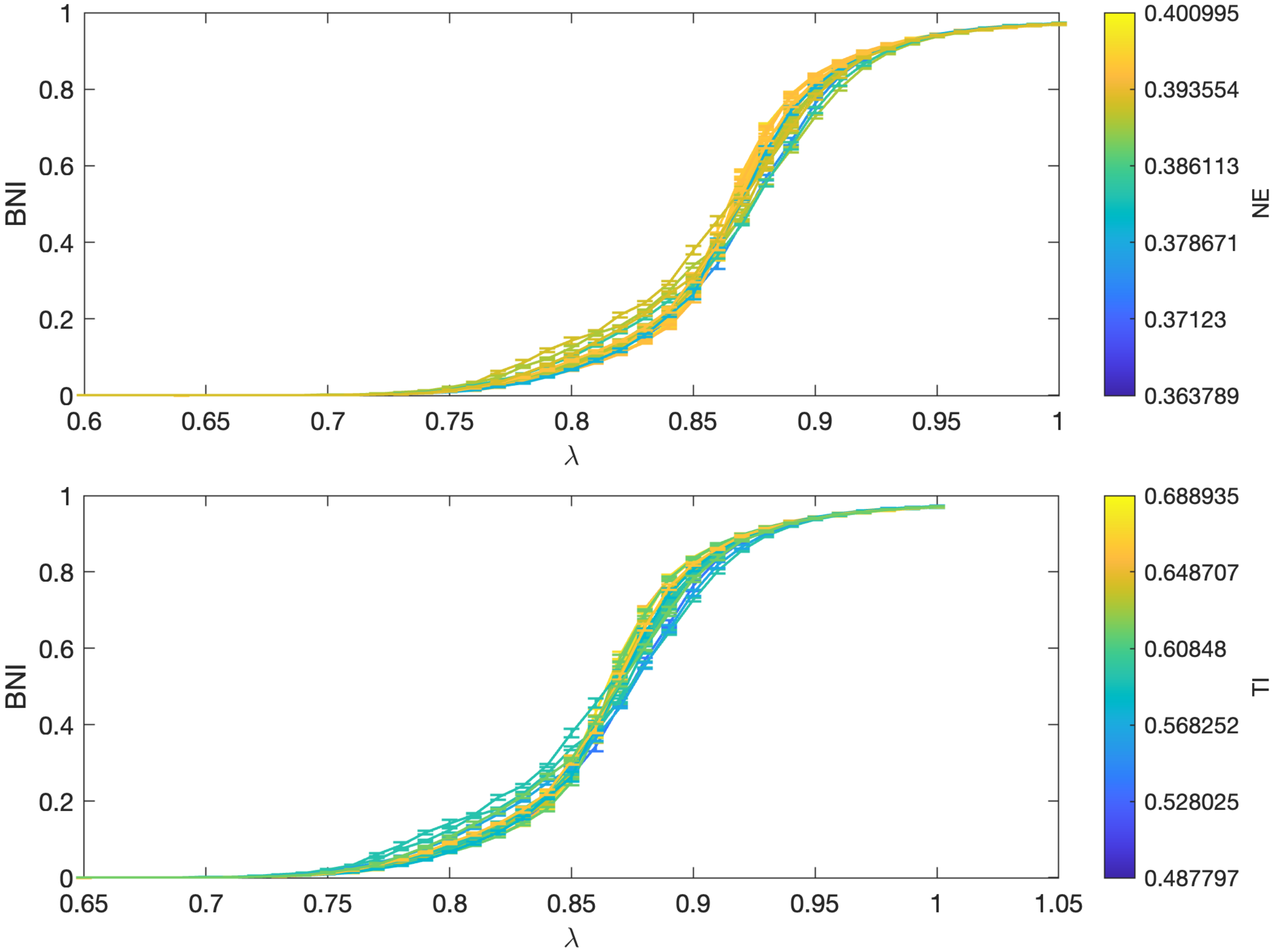


Figure S4: Increase of BNI as a function of $\lambda_0$ for 20 node networks with size of FTC, $n = 20$ and a mean degree of 2.5. Network trajectories are coloured according to their efficiency (top) and trophic incoherence (bottom).

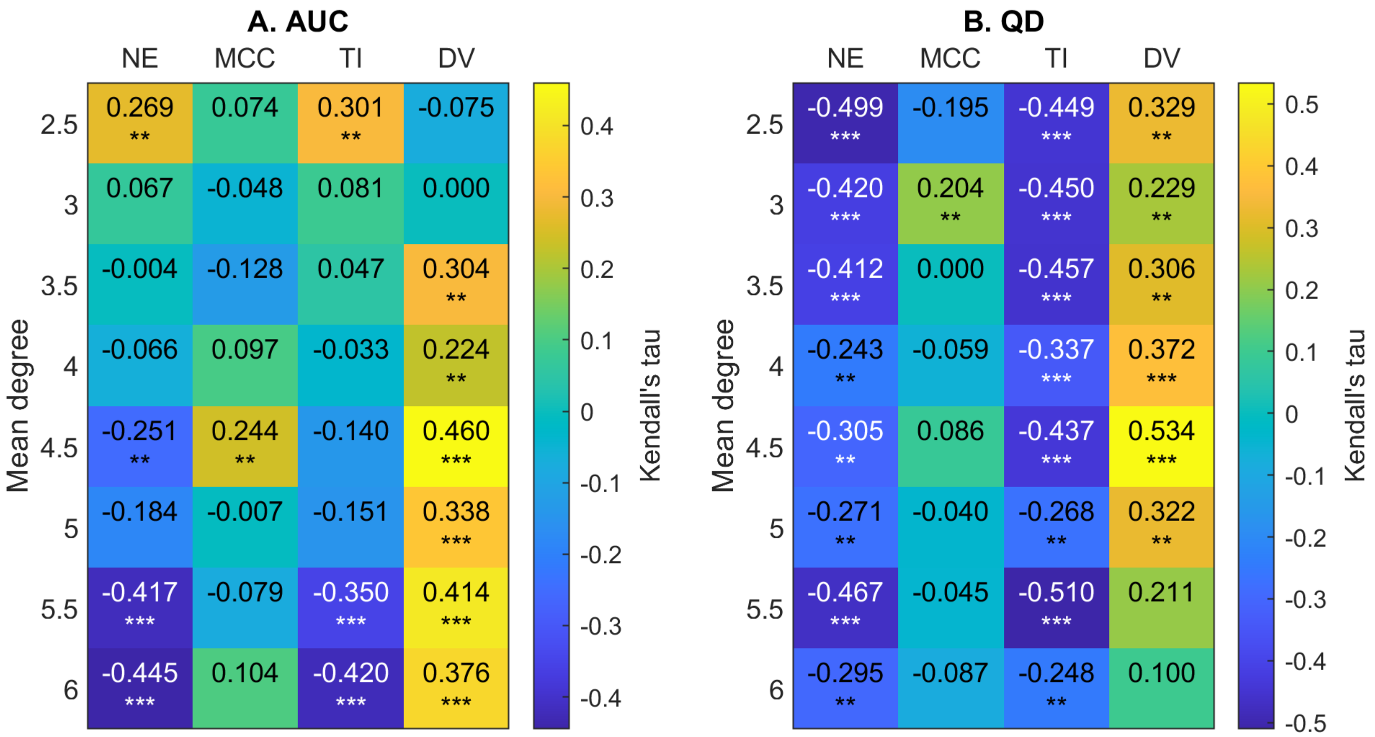


Figure S5: Heat map of Kendall correlation coefficients between network metrics (Efficiency, mean clustering coefficient, trophic incoherence and degree variance) and AUC, QD for 20 node networks generated with a increasing mean degree. A set of 50 networks with size of FTC, $n = 50$ were analysed for each value of mean degree. Significance values are labelled as * where $p < 0.05$, ** where $p < 0.01$ and *** where $p < 0.0001$.
